# Exonuclease Xrn1 regulates TORC1 signaling in response to SAM availability

**DOI:** 10.1101/2023.09.28.559955

**Authors:** Madeline M. McGinnis, Benjamin M. Sutter, Samira Jahangiri, Benjamin P. Tu

## Abstract

Autophagy is a conserved process of cellular self-digestion that promotes survival during nutrient stress. In yeast, methionine starvation is sufficient to induce autophagy. One pathway of autophagy induction is governed by the SEACIT complex, which regulates TORC1 activity in response to amino acids through the Rag GTPases Gtr1 and Gtr2. However, the precise mechanism by which SEACIT senses amino acids and regulates TORC1 signaling remains incompletely understood. Here, we identify the conserved 5’-3’ RNA exonuclease Xrn1 as a surprising and novel regulator of TORC1 activity in response to methionine starvation. This role of Xrn1 is dependent on its catalytic activity, but not on degradation of any specific class of mRNAs. Instead, Xrn1 modulates the nucleotide-binding state of the Gtr1/2 complex, which is critical for its interaction with and activation of TORC1. This work identifies a critical role for Xrn1 in nutrient sensing and growth control that extends beyond its canonical housekeeping function in RNA degradation and indicates an avenue for RNA metabolism to function in amino acid signaling into TORC1.

## Introduction

Autophagy is a conserved process that degrades cytoplasmic components and organelles under nutrient stress to maintain cell viability. The TORC1 kinase complex is a master regulator of autophagy (and many other cellular processes) in response to nutrient cues such as the availability of amino acids. The activation of TORC1 promotes cell growth and proliferation, while inhibition of TORC1 blocks cell growth and induces autophagy. Precisely how cells sense a variety of nutrient cues and their metabolic state to regulate TORC1 activity and autophagy has yet to be fully elucidated.

When budding yeast cells (*S. cerevisiae*) are switched from a nutrient-rich, lactate-based media (YPL) to a synthetic, minimal, lactate media (SL), they induce autophagy despite abundant nitrogen^1^. This autophagy-inducing regime, which forces yeast cells to utilize mitochondrial respiration, has allowed for the discovery of additional regulators of autophagy such as the SEACIT complex^1–3^, called GATOR1 in mammalian cells^4^, which senses amino acid availability. As autophagy under these conditions is potently suppressed by the sole addition of methionine, SEACIT can be seen as a regulator of autophagy triggered by an insufficiency of methionine and its downstream metabolite S-adenosyl methionine (SAM)^2^.

SEACIT has been found to sense cellular state and inhibit TORC1 through mechanisms at the post-translational level^5,6^. However, several recent studies suggest autophagy can also be regulated at the post-transcriptional level. The RNA exonuclease Xrn1 and decapping enzyme Dcp2 have been implicated in the regulation of autophagy under nitrogen starvation through a mechanism involving RNA decay^7–9^. Xrn1 is an evolutionarily conserved cytoplasmic 5’-3’ exonuclease responsible for the degradation and processing of wide variety of RNAs, including mRNAs, tRNAs, rRNAs, and lncRNAs^10–13^. RNA decay occurs simultaneously at both the 5’ and 3’ ends. mRNAs are first trimmed and/or deadenylated at the 3’ end by the Pan2-Pan3 and CCR4-NOT complexes, and then undergo decapping, removal of the 5’ 7-methylguanosine cap by Dcp2, before being degraded by Xrn1.

The reported mechanisms of post-transcriptional autophagy regulation involve the sequestration and degradation of autophagy (*ATG*) mRNA transcripts under nutrient-replete conditions, thus suppressing autophagy. Nitrogen starvation reduces levels of Xrn1 protein, resulting in accumulation of *ATG* gene transcripts, promoting autophagy induction^8^. RCK family RNA helicases Dhh1, Vad1, and DDX6 and 3’ RNA decay complex Pat1-Lsm have also been implicated in regulation of autophagy under nitrogen starvation through similar mechanisms^7,9^. However, how these RNA helicases and decay enzymes achieve specificity for *ATG* gene transcripts over other mRNAs is unclear.

We sought to further understand the role of Xrn1 in the regulation of autophagy. We find that Xrn1 has a prominent role in the regulation of autophagy induced by methionine starvation. In particular, cells lacking Xrn1 exhibit increased inhibition of TORC1 and enhanced autophagy. Unexpectedly, we found that Xrn1 regulates autophagy in a manner that is independent of *ATG* mRNA accumulation. Instead, Xrn1 exhibits biochemical and genetic interactions with the SEACIT complex and regulates TORC1 activity through toggling the nucleotide binding-state of the small Rag GTPase proteins Gtr1 and Gtr2. These findings reveal a fundamental role for this RNA exonuclease in the direct regulation of TORC1 signaling in response to amino acids and suggest a mechanism for signaling between RNA decay and cell growth control in response to nutritional stress.

## Results

### Cells lacking Xrn1 exhibit enhanced autophagy and are unresponsive to methionine in respiratory conditions

Xrn1 has been implicated in the regulation of autophagy during nitrogen starvation^8^, but a role for this exonuclease in regulating autophagy in response to methionine deprivation has not been determined. To assess the role of Xrn1 under such conditions, we grew wild type (WT) and Xrn1 null yeast strains (*xrn1Δ*) in a rich, lactate-based media (YPL), then switched cells to a synthetic, minimal, lactate media (SL) (Figure 1A). This media switch robustly induces autophagy^1,2^. We observed that *xrn1Δ* cells induced autophagy more readily than WT, even in YPL media when autophagy is normally repressed (Figures 1B-C).

**Fig. 1.**
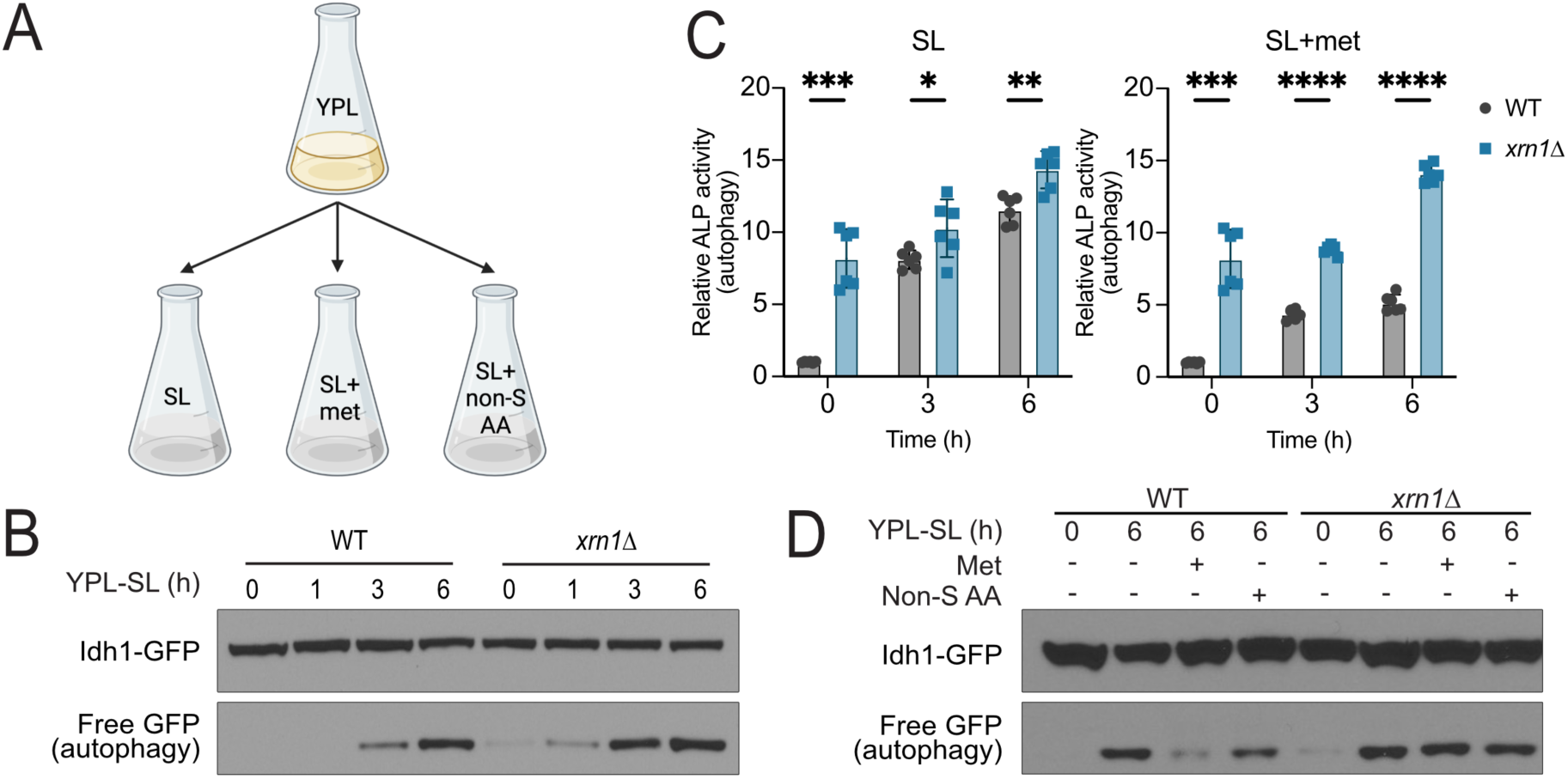
Xrn1 is a negative regulator of autophagy induced by methionine deprivation. **(A)** Scheme depicting the media switch from rich lactate medium (YPL) to minimal, synthetic lactate media (SL). Cells are supplemented with either 1 mM methionine (met) or non-sulfur amino (non-S AA) acid mix containing 1 mM of all amino acids except methionine, cysteine, and tyrosine. **(B)** Cells lacking Xrn1 exhibit increased autophagy. Autophagy was measured by the Idh1-GFP cleavage assay^34^. The accumulation of free GFP following switch to SL medium indicates autophagy. Note *xrn1Δ* cells induce autophagy even in rich YPL media. **(C)** Cells lacking Xrn1 exhibit increased autophagy, which cannot be inhibited by methionine. Autophagy was assayed quantitatively by the alkaline phosphatase (ALP) activity assay. WT and *xrn1Δ* cells were switched from YPL to SL for the indicated times, in the absence or presence of methionine supplementation. Note autophagy is inhibited by the addition of methionine to WT but not *xrn1Δ* cells. Mean±SD, n=6, statistical analysis performed using unpaired student’s t-test. **(D)** Methionine is unable to repress autophagy in cells lacking Xrn1. As shown by the GFP cleavage assay, autophagy is inhibited by addition of methionine but not by a mix of non-sulfur amino acids in WT cells, but not *xrn1Δ* cells.

The switch from YPL media to SL media induces autophagy specifically in response to insufficiency of methionine. If cells are switched to SL and supplemented with methionine, autophagy is repressed, while the addition of all other non-sulfur amino acids has minimal effect^2^. We found that *xrn1Δ* cells are unable to repress autophagy despite methionine supplementation, thus revealing a previously unknown role for Xrn1 in methionine sensing (Figures 1C,D). Importantly, the amount of Xrn1 protein did not change in response to nutrient availability (Figures S1A,B). Moreover, the addition of methionine to SL media restored growth of WT but not *xrn1Δ* cells, consistent with the inability of methionine to inhibit autophagy in this mutant (Figure S1C). These results indicate that Xrn1 is a negative regulator of autophagy induced in response to methionine deprivation.

Switch from YPL to SL results in a significant decrease in intracellular levels of methionine and SAM, consistent with autophagy induction due to methionine insufficiency^14^. Methionine levels were comparable between *xrn1Δ* and WT cells following switch to SL media, though many other sulfur-containing metabolites, including SAM, were unexpectedly elevated in *xrn1Δ* cells (Figures S1D,E, Table S1). As such, the amplified induction of autophagy exhibited by *xrn1Δ* cells cannot be explained by insufficient methionine or other sulfur-containing metabolites. Instead, they exhibit hallmarks of enhanced but dysregulated sulfur metabolism.

As previously reported, we also observed enhanced autophagy induction in *xrn1Δ* cells following nitrogen starvation in high glucose conditions (YPDèSD-N)^8^ (Figures S2A, B). However, since the difference in autophagy between *xrn1Δ* and WT cells was more significant under methionine deprivation, and methionine did not inhibit autophagy in *xrn1Δ* mutant cells, we focused on investigating the role of Xrn1 in the regulation of autophagy in response to methionine starvation (YPLèSL).

### Cells lacking Xrn1 accumulate transcripts of core autophagy genes

A major function of Xrn1 in cells is to degrade mRNA from the 5’ end following the removal of the 5’ cap. To determine if its exonuclease activity is required for its regulation of autophagy, we generated cells expressing a catalytically dead mutant of Xrn1 (D208A)^15^. The autophagy phenotype of these cells matched that of *xrn1Δ* cells, (Figure 2A), demonstrating that Xrn1 exonuclease activity is required for normal autophagy regulation. Both the Xrn1 D208A and *xrn1Δ* mutants exhibited a severe growth defect (Figure 2B).

**Fig. 2.**
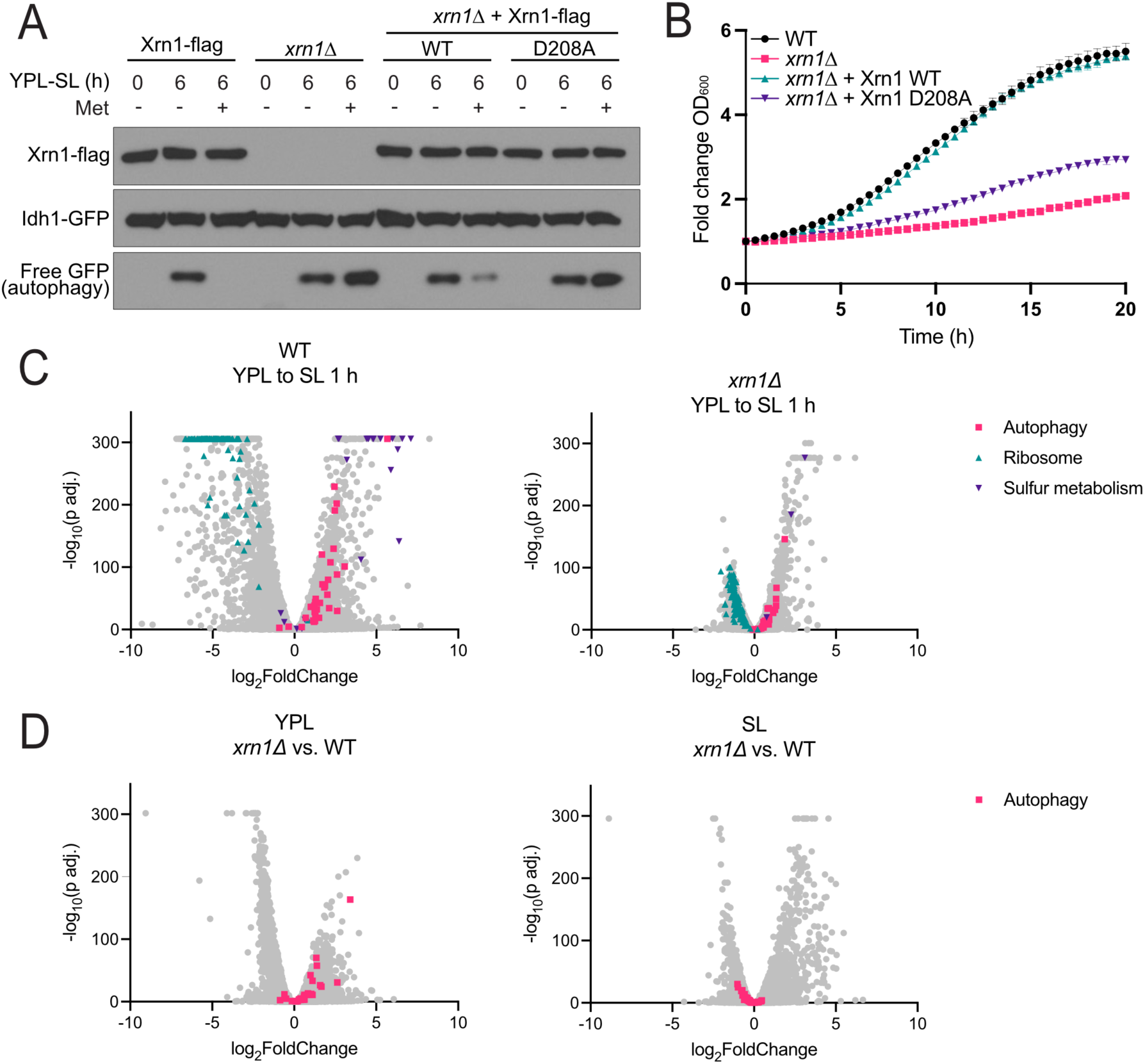
The regulation of autophagy by Xrn1 is dependent on its catalytic activity, but not dysregulation of autophagy-related mRNA transcripts. **(A)** Catalytic activity of Xrn1 is required for methionine-responsive autophagy regulation. Xrn1 was knocked out and WT or catalytically dead (D208A) flag-tagged Xrn1 was expressed ectopically from a plasmid using the endogenous Xrn1 promoter. WT Xrn1, but not Xrn1 D208A, was able to rescue the methionine-sensitive autophagy phenotype of *xrn1Δ* cells. **(B)** Cells lacking Xrn1 exonuclease activity have a severe growth defect. The indicated strains were grown in YPL media and growth rate was monitored by automated measurement of OD_600_ every 30 min. **(C)** Differentially expressed genes in WT and *xrn1Δ* cells before and after the switch to SL for 1 h. Known differentially expressed gene groups are highlighted to demonstrate their muted response in *xrn1Δ* compared to WT cells. **(D)** Differentially expressed genes between *xrn1Δ* and WT cells in either YPL or SL media. Autophagy (*ATG*) genes are highlighted. RNA-seq data are also available in Table S2.

Xrn1 plays a general role in regulating gene expression through RNA decay. It has been reported that Xrn1 regulates the levels of core autophagy (*ATG*) gene mRNAs in response to nitrogen availability, allowing for accumulation of these genes under starvation to induce autophagy^8^. We examined whether the loss of Xrn1 affected levels of *ATG* mRNAs under methionine deprivation by RNA-seq. WT cells induce sulfur metabolism (e.g., *MET*) genes and repress growth-promoting genes (e.g., ribosomal subunits) following the switch to SL, consistent with methionine starvation^16^. In contrast, *xrn1Δ* cells displayed fewer overall changes in transcript abundance following the switch from YPL to SL (Figure 2C, Table S2). The abundance of *ATG* mRNAs was increased in cells lacking Xrn1 in YPL media, but was decreased in SL media (Figures 2D, S3B). As autophagy is upregulated in *xrn1Δ* cells compared to WT cells in this condition, these transcriptomic analyses reveal a disconnect between the abundance of *ATG* mRNA transcripts and the induction of autophagy. Collectively, these results suggest that alteration of specific *ATG* transcripts is not the primary mechanism by which Xrn1 regulates autophagy induced by methionine starvation.

### Cells lacking Xrn1 exhibit diminished TORC1 signaling

TORC1 is a master regulator of autophagy^17,18^. To assess whether Xrn1 might regulate autophagy through TORC1, we assayed TORC1 signaling by examining the phosphorylation of one of its downstream substrates, ribosomal protein S6 (Rps6). Switch to minimal media (SL) results in reduced S6 phosphorylation (S235/236), consistent with TORC1 inhibition and induction of autophagy, which can be restored by addition of methionine to minimal media (Figure 3A). Strikingly*, xrn1*Δ cells have reduced p-S6, even in YPL rich media. In SL media, p-S6 is further diminished, and addition of methionine to the media has no effect on p-S6 levels in cells lacking Xrn1 (Figure 3A). Xrn1 is therefore required for the ability of TORC1 to properly sense methionine availability. Furthermore, the amounts of p-S6 observed in *xrn1Δ* cells are inversely correlated with the level of autophagy, consistent with a potential role of Xrn1 in regulating autophagy through TORC1.

**Fig. 3.**
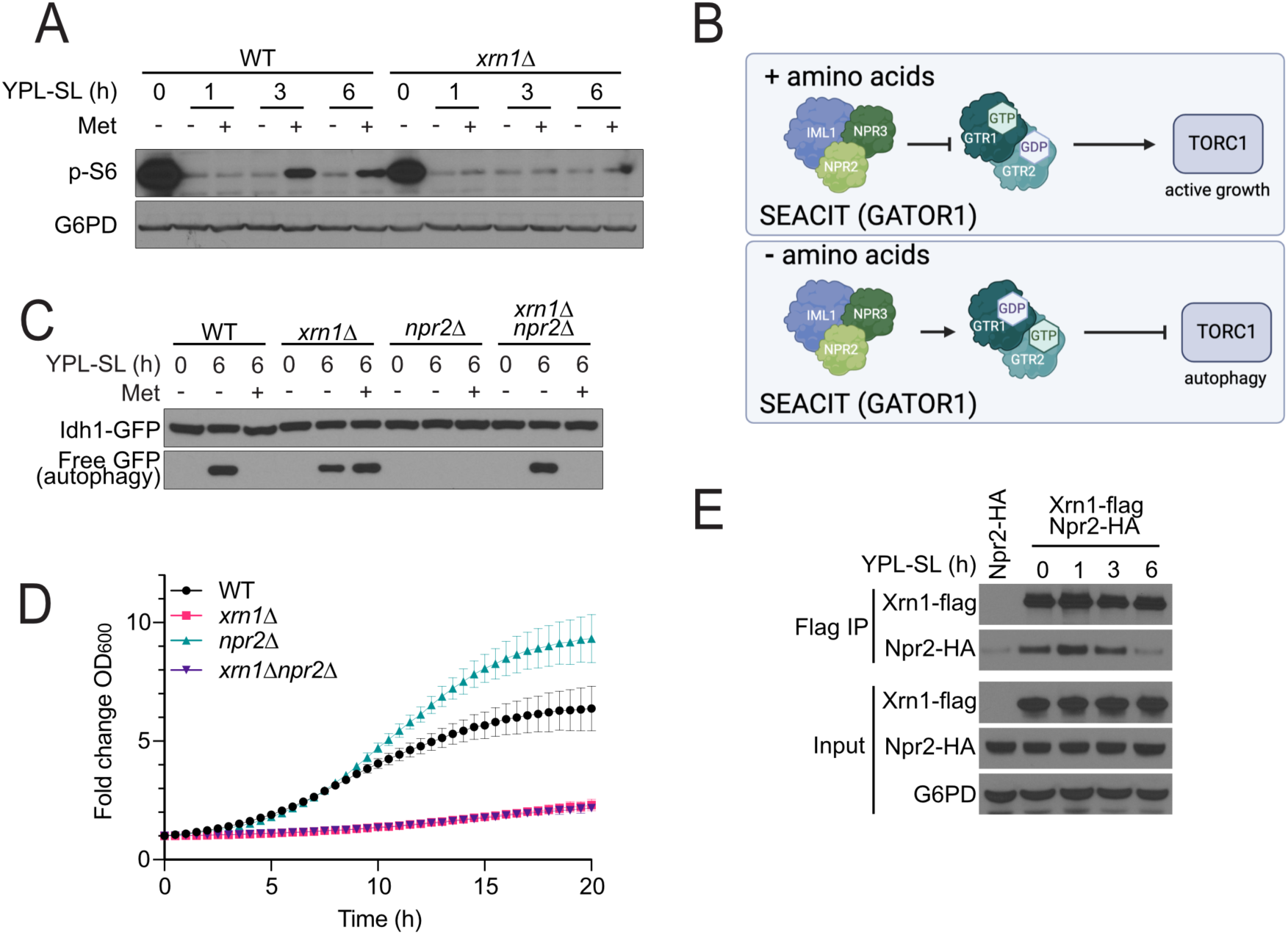
Xrn1 regulates autophagy through modulation of TORC1 signaling. **(A)** Cells lacking Xrn1 exhibit diminished TORC1 activity. WT and *xrn1Δ* cells were grown in YPL and switched to SL media, or SL containing methionine, for the indicated times. Western blot analysis was used to assess phosphorylation of the TORC1-dependent substrate, ribosomal protein S6. **(B)** Npr2 and the SEACIT complex regulate TORC1 activity through the Rag GTPases Gtr1 and Gtr2. The nucleotide binding state of this heterodimer determines whether the complex activates or inhibits TORC1. **(C)** GFP cleavage autophagy assay in cells lacking Xrn1 and/or Npr2. The indicated strains were grown in YPL and then switched to SL for 6 h, in the absence or presence of 1 mM methionine. **(D)** Growth curves of cells lacking Xrn1, Npr2, or both. The indicated strains were grown in YPL and OD_600_ was measured every 30 min. Fold change in OD_600_ is plotted. **(E)** Xrn1 interacts with Npr2. Cells expressing Flag-tagged Xrn1 and HA-tagged Npr2 were grown in YPL and then switched to SL for the indicated times. Cells were harvested and the interaction between Xrn1 and Npr2 was determined by co-immunoprecipitation followed by Western blotting.

### Xrn1 interacts with the SEACIT complex, a negative regulator of TORC1

We next sought to understand the mechanism by which Xrn1 might regulate TORC1. The SEACIT complex (comprised of Npr2, Npr3, Iml1) is a key regulator of TORC1 activity through the Rag GTPase heterodimer Gtr1 and Gtr2 in response to amino acid levels. The role of Gtr1/2 in regulating TORC1 is dependent on their nucleotide-binding state. When amino acids are sufficient, Gtr1 is loaded with GTP, while Gtr2 is bound to GDP^19,20^. In this state, the Gtr1/2 complex activates TORC1 and promotes cell growth^21^. The SEACIT complex is a dedicated GTPase activating protein (GAP) for Gtr1^22^. When amino acids are insufficient, SEACIT triggers the hydrolysis of the GTP bound by Gtr1, and the exchange of GDP for GTP on Gtr2, thereby inactivating the Gtr1/2 complex, which in turn leads to TORC1 inhibition and autophagy induction^19,20^ (Figure 3B). Deletion of any component of SEACIT renders TORC1 constitutively active and inhibits autophagy^1,3,21–23^.

The SEACIT component Npr2 has been found to be selectively required for autophagy induction under respiratory conditions, but not nitrogen starvation^1^. We examined the regulation of autophagy in cells lacking both Xrn1 and Npr2. Unexpectedly, the loss of both proteins restored normal, methionine-responsive regulation of autophagy (Figure 3C). This result suggests that Xrn1 may function in this pathway to control TORC1 activity, and that these two proteins may have opposing roles in TORC1 regulation.

To further understand the role of Xrn1 in relation to the SEACIT complex, we examined the growth of cells lacking Xrn1, Npr2, or both. Loss of Npr2 results in increased growth under respiratory conditions, consistent with the role of Npr2 in TORC1 inhibition (Figure 3D). Cells lacking Xrn1 have a severe growth defect, and the further loss of Npr2 has no effect on growth, indicating that Xrn1 may function downstream of Npr2, but upstream of TORC1.

We next tested whether Xrn1 might associate with Npr2 or other components of the SEACIT complex. Co-immunoprecipitation (co-IP) analysis revealed that Xrn1 interacts with Npr2 in YPL media, and that interaction is decreased following switch to SL media (Figure 3E), suggesting their interaction is dependent on methionine availability.

### Xrn1 regulates Gtr1/2 nucleotide-binding state

Our results indicate that Xrn1 regulates autophagy by controlling TORC1 activity downstream of Npr2 and the SEACIT complex. Since SEACIT is a GAP for Gtr1, we next assayed the methionine sensitivity of cells lacking Gtr1 alone or in combination with Xrn1. The loss of Gtr1 alone did not result in changes in autophagy. However, in combination with the loss of Xrn1, these cells became entirely insensitive to methionine availability, inducing autophagy in all media conditions (Figure 4A). Furthermore, we observed that Xrn1 interacts with Gtr1 independent of methionine starvation (Figure S4A).

**Fig. 4.**
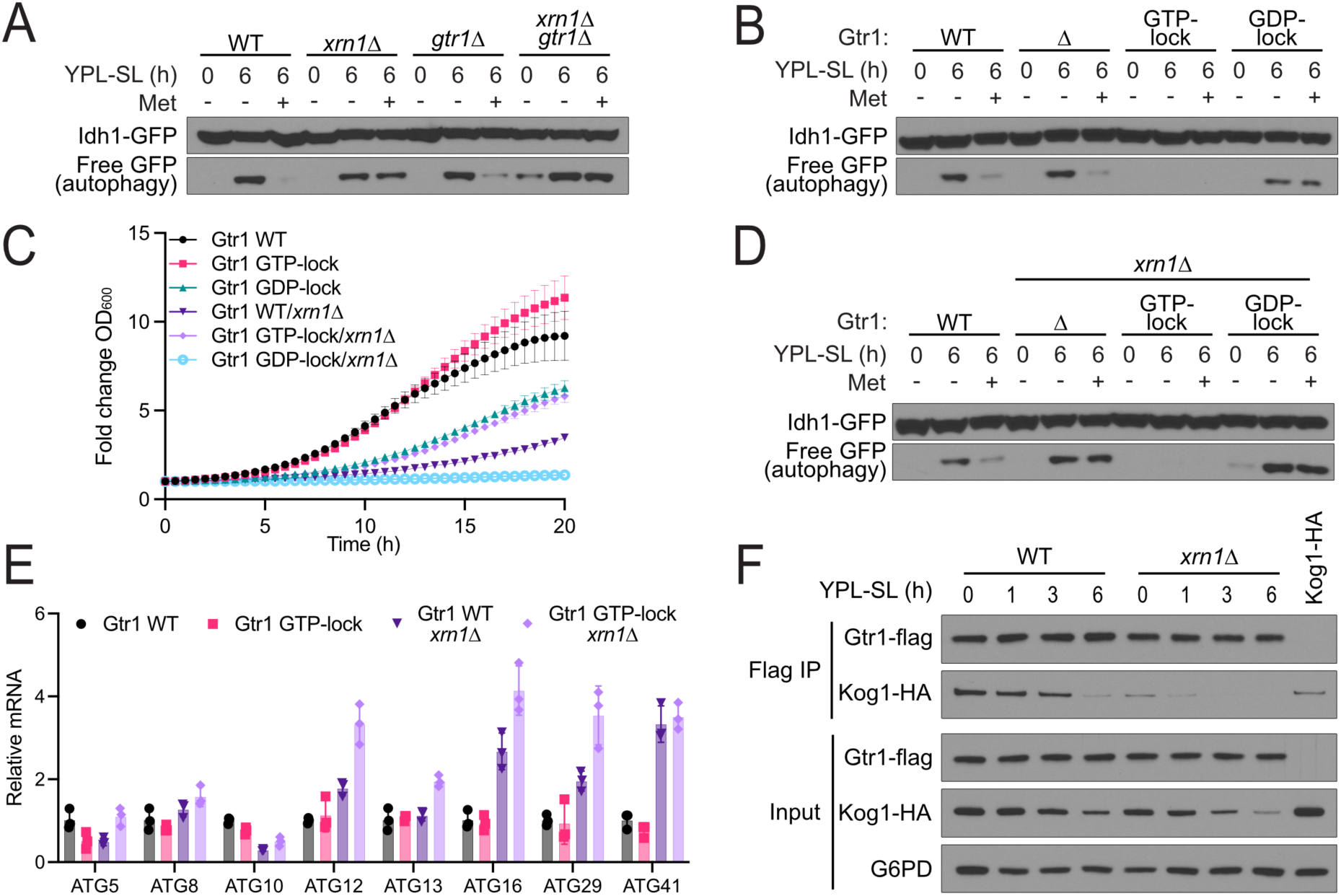
Xrn1 regulates autophagy through the Rag GTPase proteins Gtr1/2. **(A)** The loss of Xrn1 and Gtr1 results in autophagy induction regardless of methionine availability. The indicated strains were grown in YPL and then switched to SL for 6 h, in the absence or presence of 1 mM methionine. Autophagy was measured using the GFP-cleavage assay. **(B)** The nucleotide-binding state of Gtr1 regulates autophagy. Point mutations of Gtr1 to lock it in its GTP-binding (Q65L) or GDP-binding (S20L) state affect its role in regulating autophagy through TORC1. Autophagy was assayed as in (A). **(C)** GTP-and GDP-locked mutations in Gtr1 predominate the growth defect of *xrn1Δ* cells. The indicated strains were grown in YPL and OD_600_ was measured every 30 min. Fold change in OD_600_ is plotted. **(D) (E)** GTP-and GDP-locked mutations in Gtr1 are sufficient to bypass the role of Xrn1 in regulating autophagy following methionine deprivation. The indicated strains were grown in YPL and then switched to SL for 6 h, in the absence or presence of 1 mM methionine. Autophagy was measured as in (A). **(F)** GTP-locked mutation of Gtr1 is unable to restore WT levels of many autophagy gene transcripts despite restoration of TORC1 signaling, repression of autophagy, and enhanced growth. mRNA abundance was assayed by qPCR for the indicated genes. **(G)** Loss of Xrn1 results in decreased interaction between Gtr1 and TORC1 component Kog1. WT and *xrn1Δ* cells expressing Gtr1-Flag and Kog1-HA were grown in YPL and switched to SL for the indicated times. The interaction between Gtr1 and Kog1 was assessed by co-IP followed by Western blotting.

Specific mutations in Gtr1 confer dominant regulation of TORC1 activity and autophagy^20,21,24,25^. Cells expressing GTP-locked Gtr1 (Q65L) are unable to induce autophagy following switch to SL media and exhibit enhanced growth (Figures 4B,C). Conversely, cells expressing GDP-locked Gtr1 (S20L) induce autophagy regardless of methionine availability and have a slow-growth phenotype (Figures 4B,C). Strikingly, when these mutations were introduced in the *xrn1Δ* background, their phenotypes became dominant to *xrn1Δ*: cells with GTP-locked Gtr1 cannot induce autophagy and have enhanced growth, while cells with GDP-locked Gtr1 constitutively induce autophagy and have a severe growth defect (Figure 4C, D). The GTP-locked mutation of Gtr1 is also sufficient to restore TORC1 kinase activity in cells lacking Xrn1 (Figure S4B). Moreover, Xrn1 preferentially interacted with the GDP-locked form of Gtr1, indicating that Xrn1 may have a specific role in TORC1 regulation through Gtr1 (Figure S4C).

Notably, although loss of Xrn1 can cause accumulation of *ATG* mRNA transcripts, we find that the mutation of Gtr1 to its GTP-locked state does not restore WT levels of these transcripts (Figure 4E). This result indicates that, although this mutant is dominant in its suppression of autophagy through constitutive activation of TORC1, cells with this mutation still erroneously accumulate mRNAs of key autophagy genes. Collectively, these results decouple the abundance of autophagy mRNAs from the induction of autophagy, and supports a model in which Xrn1 regulates autophagy through modulation of TORC1 signaling directly, rather than through degradation of key mRNAs.

Consistent with previous reports, mutations in Gtr2 have a milder effect on autophagy regulation, yet these mutations, as well as loss of Gtr2, exhibit synergistic effects in combination with the loss of Xrn1 (Figures S5A,B,C)^21,25^. Thus, mutations that alter the nucleotide-binding state of Gtr1 are epistatic to *xrn1Δ*, while mutations that alter the nucleotide-binding state of Gtr2 synergize with *xrn1Δ* in terms of autophagy phenotypes. These data are consistent with a model in which Xrn1 converts Gtr1/2 from a TORC1-inhibitory state to a TORC1-activating state.

Gtr1 has one annotated guanine nucleotide exchange factor (GEF), Vam6^20^. Gtr2 has two GAPs, Lst4 and Lst7, which act in a complex (Figure S5D)^26,27^. We tested autophagy induction in response to methionine starvation of strains lacking these proteins alone or in combination with *xrn1Δ.* Like *xrn1Δ* mutants, cells lacking Lst4 or Lst7 lost the ability to sense methionine and induced autophagy regardless of its availability (Figures S5E,F). Moreover, the loss of Xrn1 in addition to either Lst4 or Lst7 severely exacerbated the methionine-insensitive autophagy and growth defect phenotypes of these strains (Figures S5E,G). These genetic epistasis experiments strongly suggest that Xrn1 functions by toggling Gtr1/2 nucleotide binding states in a manner conducive for TORC1 activation, but might not act through Lst4 and Lst7. Unexpectedly, cells lacking Vam6 were unable to induce autophagy following methionine deprivation, suggesting that the role of this protein in respiratory conditions is not fully understood (Figure S5H). Ivy1 and Ait1 have been described as regulators of TORC1 signaling through Gtr1 and Gtr2^28,29^. Loss of either of these proteins had no effect on autophagy regulation in response to methionine deprivation (Figure S5I).

### Xrn1 is required for Gtr1/2 regulation of TORC1 activity

As the GTP-bound form of Gtr1 interacts with TORC1^20,21,25^, we next assessed the interaction between Gtr1 and Kog1, a component of TORC1, in *xrn1Δ* cells. Strikingly, the interaction between Gtr1 and Kog1 was significantly reduced even in the presence of methionine, consistent with reduced TORC1 activity and enhanced autophagy in *xrn1Δ* cells (Figure 4F). However, we also noted that Kog1 exhibits markedly reduced solubility in cells lacking Xrn1 and in WT cells following switch to methionine deprivation (Figure S5J), consistent with reports of inactivated TORC1 re-localization to vacuolar domains^30^. Collectively, our results strongly suggest that Xrn1 promotes the GTP-bound state of Gtr1 to enable TORC1 activation.

### Xrn1 interactions with Npr2 and Gtr1 are dependent on RNA availability

Because the catalytic activity of Xrn1 is required for its role in autophagy regulation, we next tested the catalytically-inactive D208A mutant and found that it exhibited reduced interactions with Npr2 and, to a lesser extent, Gtr1 (Figures 5A,B). Consistent with these observations, the addition of RNase A to the co-IP buffer enhanced the interaction of Xrn1 with both Npr2 and Gtr1, suggesting that these protein interactions can be sensitive to RNA (Figures 5C,D).

**Fig. 5.**
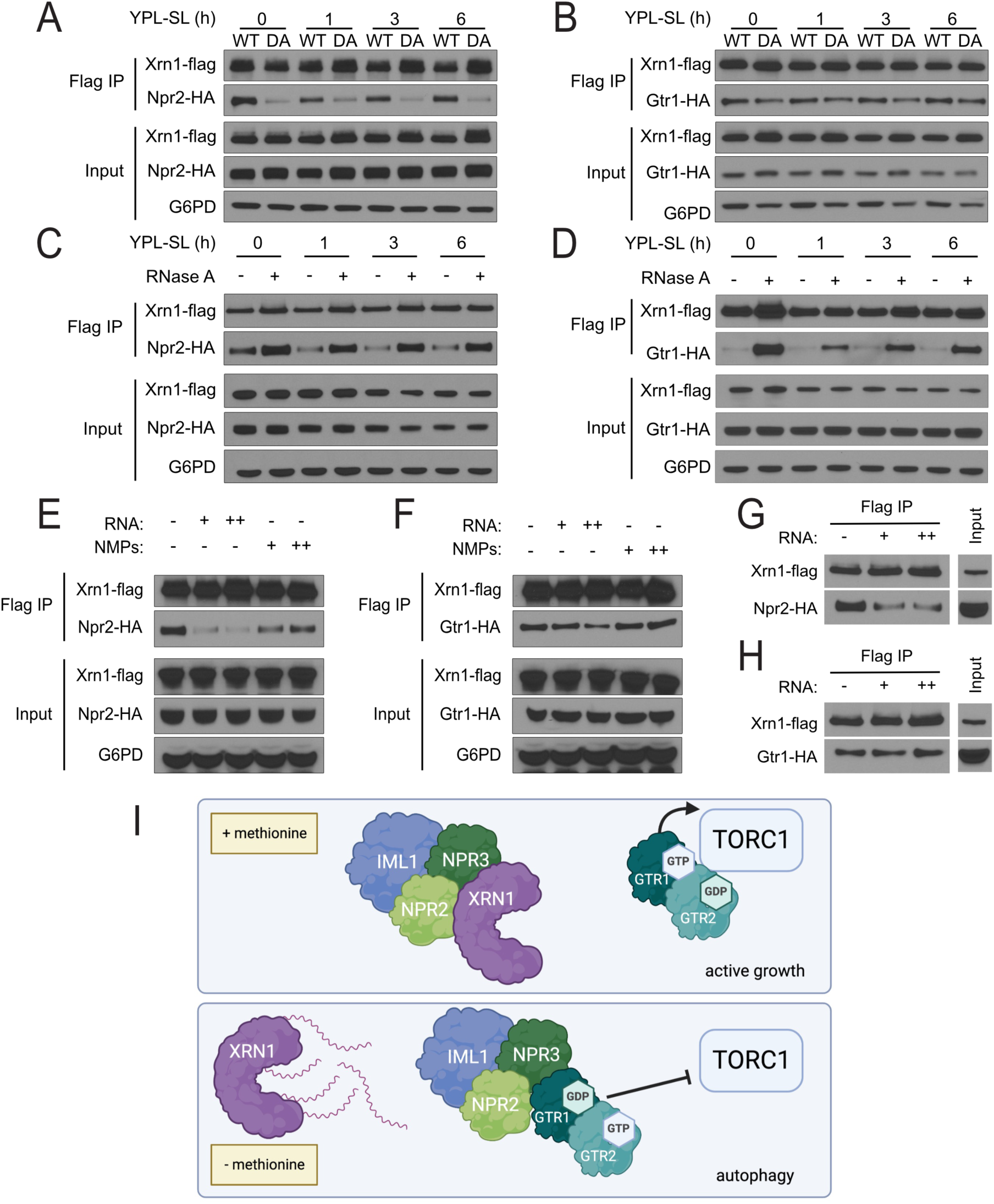
The interaction between Xrn1 and Npr2 is regulated by RNA availability. **(A), (B)** Interaction between catalytically dead Xrn1 mutant and Npr2 or Gtr1. In *xrn1Δ* cells expressing either HA-tagged Npr2 (A) or Gtr1 (B), WT or catalytically dead (D208A = DA) flag-tagged Xrn1 was expressed ectopically from a plasmid using the endogenous Xrn1 promoter. Cells were grown in the indicated conditions, and the interaction between Xrn1 and Npr2 or Gtr1 was assessed by co-IP followed by Western blotting. **(C), (D)** The interaction of Xrn1 with Npr2 (C) or Gtr1 (D) is sensitive to RNase treatment. Cells with flag-tagged Xrn1 and either HA-tagged Npr2 or Gtr1 were grown in the indicated conditions. The interaction between Xrn1 and Npr2 or Gtr1 was assessed by co-IP followed by Western blotting. Addition of RNase A (25 ng/μL) to the co-IP lysate increases interaction of Xrn1 with both Npr2 and Gtr1. **(E), (F)** The interaction of Xrn1 with Npr2 (E) but not Gtr1 (F) is sensitive to RNA concentration. Cells with flag-tagged Xrn1 and either HA-tagged Npr2 or Gtr1 were grown in YPL. The interaction between Xrn1 and Npr2 or Gtr1 was assessed by co-IP with RNA or NMPs added to the lysate followed by Western blotting. The addition of total RNA (from exponentially growing WT cells in YPL media) at 125 μg/mL (+) or 313 μg/mL (++), but not free NMPs, is sufficient to disrupt the interaction between Xrn1 and Npr2, but not Xrn1 and Gtr1. **(G), (H)** Treatment with RNA was sufficient to disrupt the interaction of Xrn1 with Npr2 (G) but not Gtr1 (H). Cells with flag-tagged Xrn1 and either HA-tagged Npr2 or Gtr2 were grown in YPL. The interaction between Xrn1 and Npr2 or Gtr1 was assessed by co-IP, followed by addition of RNA to the wash buffer, then analyzed by Western blotting. **(I)** Model for regulation of TORC1 activity by Xrn1 through Gtr1/2.

Finally, we investigated whether the presence of RNA might influence interactions between Xrn1 and either Npr2 or Gtr1. The addition of total RNA isolated from cells grown in YPL, either during or after the co-IP, was sufficient to reduce the interaction between Xrn1 and Npr2, but not Xrn1 and Gtr1 (Figure 5E-H). As a comparison, the addition of nucleotide monophosphates (NMPs), which are the products of RNA degradation by Xrn1, had no effect on Xrn1 interactions as assessed by co-IP. Collectively, these results indicate that cellular RNA levels may regulate the interaction of Xrn1 with SEACIT, allowing for differential regulation of TORC1 activity in response to local or global RNA accumulation in cells.

## Discussion

In this work, we have identified a role for an RNA exonuclease, Xrn1, in TORC1 signaling through regulation of Gtr1/2 nucleotide-binding states. Loss of Xrn1 results in reduced interaction between Gtr1 and TORC1, which inhibits the complex^25^. Curiously, cells lacking Xrn1 also exhibit a severe growth defect, consistent with previous observations^31^. We have shown here that this defect can be largely rescued by a single mutation of Gtr1 locking it in its GTP-bound conformation. Thus, the severe growth defect of *xrn1Δ* cells cannot be solely attributed to a defect in RNA decay, but is instead due to TORC1 inhibition.

Prior studies have linked RNA decay enzymes to the regulation of autophagy through the expression of key autophagy-driving mRNAs^7–9^. In response to methionine starvation, we observed minimal changes in mRNA levels of the majority of *ATG* genes in the absence of Xrn1. Instead, our findings suggest a direct role for an RNA decay enzyme (i.e., Xrn1) in the regulation of autophagy through TORC1 under conditions of nutritional stress.

How might Xrn1 sense methionine availability to regulate TORC1 activity? Methionine starvation triggers significant changes in the transcriptome (Figure 2C). Transcripts coding for proteins involved in growth and ribosome biogenesis are rapidly degraded, while genes for sulfur metabolism must be transcribed to promote methionine and SAM biosynthesis for survival^16^. Moreover, methionine starvation leads to an insufficiency of SAM, which could result in compromised ability to methylate particular RNAs, including tRNAs and rRNAs. Xrn1 has been shown to have not only a general role in buffering cellular RNA levels that cannot be managed by other RNA decay enzymes^32^, but also a specific role in degrading particular RNAs^13,33^. We report here that Xrn1 is capable of interactions with Npr2, which can be influenced by the presence of RNA. The accumulation of RNAs that need to be degraded in response to an insufficiency of SAM may titrate Xrn1 away from SEACIT, thereby liberating SEACIT to act on Gtr1, converting it to its GDP-bound state and inhibiting TORC1. In cells lacking Xrn1, the SEACIT complex is free to act on Gtr1, shifting the equilibrium towards the inactive state of this complex and promoting autophagy (Figure 5I).

As for how precisely Xrn1 regulates Gtr1/2, our cumulative findings suggest that Xrn1 likely acts by regulating the nucleotide binding states of the Gtr1/2 heterodimer. Notably, GTP-locked Gtr1 mutants bypass *xrn1Δ* phenotypes, while in stark contrast GTP-locked Gtr2 mutants exacerbate them. As such, we hypothesize that Xrn1 likely acts as either a GEF for Gtr1 or a GAP for Gtr2, such that the loss of Xrn1 keeps the heterodimer in its inactive state, thereby inhibiting TORC1. Alternatively, Xrn1 could act through the other GAPs and GEFs that modulate Gtr1/2.

In the absence of Xrn1, our metabolite profiling experiments reveal elevated levels of many sulfur-containing metabolites, including SAM, cystathionine, and glutathione, despite reduced TORC1 signaling and enhanced autophagy. As *MET* gene transcripts were not elevated in *xrn1Δ* mutant cells, how Xrn1 contributes to sulfur metabolic processes is unclear, but is nonetheless consistent with a key role for this enzyme in the cellular response to methionine availability. Moreover, it has not escaped our attention that bulk intracellular levels of GTP and GDP were also modestly decreased in *xrn1Δ* cells (Figure S6F), suggesting that Xrn1 may also play a role in recycling of guanosine nucleotides from RNA that may be supplied for Gtr1/2 function. However, our attempts to rescue the phenotypes of *xrn1Δ* with guanosine supplementation have not been successful thus far. Nonetheless, it remains possible that Xrn1 could also help locally recycle GDP or GTP pools for GEF-mediated exchange of their bound nucleotides.

The majority of previous studies on TORC1 signaling have focused on how various amino acids, nutrients, or growth factors directly signal into TORC1 through a variety of protein complexes. In this study, we now show how RNA can also act as a signal to modulate TORC1 signaling through the exonuclease Xrn1. In this manner, the accumulation of particular RNAs, perhaps in response to reduced methylation potential, could act as a surrogate of the metabolic state and indicator of cellular stress, which can then be integrated into the regulation of cellular metabolic activities through TORC1. As the SEACIT complex and Xrn1 are conserved in mammalian cells, this mechanism of RNA signaling into TORC1 is likely to be conserved as well.

## Methods

### Strain growth and media conditions

All strains used in this study are listed in Table S3. The prototrophic yeast strain CEN.PK was used in all experiments. Gene deletions were made using standard PCR-based methods to amplify resistance-based cassettes with flanking sequences to replace the target gene sequence by homologous recombination. C-terminal tags were made with PCR-based methods to amplify resistance cassettes with flanking sequences. Mating, sporulation, and tetrad dissection were used to construct strains with multiple alleles whenever possible to minimize occurrence of compensatory or suppressor mutations.

### Media used in this study

YPL (1% yeast extract, 2% peptone, 2% lactate), YhPL (0.5% yeast extract, 2% peptone, 2% lactate), YPD (1% yeast extract, 2% peptone, 2% glucose), SL (0.67% yeast nitrogen base without amino acids, 2% lactate), YPD (1% yeast extract, 2% peptone, 2% glucose), SD-N (0.17% yeast nitrogen base without amino acids and ammonium sulfate). YhPL was used only for GFP cleavage autophagy assay. Growth curves were performed using 96-well plates and continuous OD_600_ measurements using a Tecan SparkControl Magellan plate reader. Yeast were grown at 30^°^C.

### Whole cell extract preparation

Trichloroacetic acid (TCA)-based protocols were used to lyse yeast cells for Western blots. Cell pellets were quenched in 15% TCA, 15% ethanol for 15 min on ice. Cell pellets were spun down and resuspended in cold 15% ethanol and lysed by beads beating. Lysate was pelleted and washed with cold 100% ethanol and resuspended in 2x SDS lysis buffer (4% SDS, 0.4 M Tris pH 7.5, 20% glycerol) and boiled for 5 min.

### Western blotting

Protein concentration was determined using Pierce BCA protein assay. 4x SDS sample loading dye (4% SDS, 0.24 M Tris pH 6.8, 2% SDS, 1% β-mercaptoethanol, 0.02% bromophenol blue) was added to samples before running. Similar amounts of protein were used per sample. Samples were separated by electrophoresis on 4-12% NuPAGE Bis-Tris gels. Proteins were transferred to a nitrocellulose membrane and blotted with the corresponding antibodies. Blocking and antibody incubation was performed in 5% dry milk in 1xTBST. Antibodies used were: anti-FLAG (Sigma F1804), anti-Rpn10 (Abcam, ab98843), anti-G6PD (Sigma, A9521), anti-HA (Cell Signaling, 3724), anti-GFP (Roche, 11814460001), anti-phospho-S6 (Ser235/236) (Cell Signaling, 2211), anti-mCherry (Chromotek, 5F8).

### ALP assay for autophagy

Strains expressing cytosolic Pho8Δ60 were subject to the ALP activity assay to measure the level of autophagy following the protocol from previously described methods^34^ with some modifications. Briefly, cell pellets were resuspended in 400 μL lysis buffer (250 mM Tris-HCl pH 9, 25 mM MgSO_4_, 1% Triton X-100, 1x EDTA-free protease inhibitor cocktail (Roche). Cells were lysed by beads beating with glass beads (Sigma). Cell debris was separated from the cell extract by centrifugation. 70 μL cell extracts were added to triplicate wells in 96-well flat bottom plates. 70 μL substrate solution (250 mM Tris-HCl pH 9, 25 mM MgSO_4_, 1% Triton, 2.7 mM p-nitrophenyl phosphate (MP Biomedicals)) was then added to each well. The plate was incubated at room temperature for 5 min, then the reaction was stopped with 140 μL stop buffer (1 M glycine, pH 11). The plates were read at 400 nm to determine the production of p-nitrophenol.

### GFP cleavage assay for autophagy

The release of free GFP from Idh1-GFP was examined to assay mitophagy based on previously described protocols^35^. Idh1 is a mitochondrial matrix protein. When mitophagy is induced, this protein accumulates in the vacuole and is degraded. Free GFP is more stable and is detected by Western blotting with anti-GFP antibody (Roche, 11814460001). GFP cleavage assays were performed at least 3 times and results from one representative experiment are shown.

### RNA extraction

Total RNA samples were prepared as previously described^36^. Briefly, frozen yeast cell pellets were thawed and washed with cold sterile water. Cell pellets were then resuspended in TES (10 mM Tris-HCl pH 7.5, 10 mM EDTA pH 8.0, and 0.5% SDS), followed by addition of an equal volume of acidic phenol (pH 4.3). Cells were lysed using glass beads on a beads-beater. Cell debris and glass beads were removed by centrifugation. The aqueous phase was transferred to a new tube and extracted with an equal volume of phenol. This process was repeated with a third extraction with chloroform. RNA was ethanol precipitated and resuspended in nuclease-free water.

### Fluorescence imaging

Images were taken under a 100X oil-immersion objective lens with a Zeiss LSM980 microscope. Cells grown under the indicated conditions were visualized quickly after mounting on slides. Images were processed using Zeiss Zen software.

### Immunoprecipitation

Immunoprecipitation was performed as previously described^6^. Briefly, cell pellets were harvested and stored at −80^°^C until cell lysis. The cell pellet was resuspended in 350 μL lysis buffer 1 (50 mM HEPES pH 7.5, 150 mM NaCl, 1 mM EDTA, 0.5% NP-40, 2x protease inhibitor cocktail (Roche), 1 mM PMSF, 1 mM sodium orthovanadate, 5 mM NaF, 10 μM leupeptin, 10 μM pepstatin A). Cells were lysed by beads beating with glass beads. The lysed cells were separated from beads by centrifugation and washed with 525 μL lysis buffer 2 (buffer 1, but devoid of NP-40). The cell extracts were combined and cleared by centrifugation. The protein concentration of the lysates was measured using Bradford assay (Bio-Rad) and adjusted to normalize all samples in 800 μL reaction volume. 30 μL of the input sample was taken and mixed with 4x SDS sample dye and denatured by boiling at 95^°^C for 5 minutes. For each co-immunoprecipitation reaction, 25 μL of Dynabeads Protein G (Life Technologies) was washed with the IP lysis buffer and incubated with 3 ug anti-Flag antibody (Sigma, M2, F1804) for 1 h at 4^°^C. The supernatant containing unbound antibody was then removed by centrifugation, and the conjugated bead-antibody was added to the cleared lysate. If indicated, ribonuclease A (Sigma) was added to the lysates (25 ng/μL). Where indicated, total RNA prepared from WT yeast grown in YPL or SL media, or nucleotide monophosphates (NMPs, adenosine monophosphate, cytidine monophosphate, guanosine monophosphate, uridine monophosphate) were added to the lysate. After incubating for 2 h at 4^°^C, the beads were washed 3 times with wash buffer (50 mM HEPES pH 7.5, 150 mM NaCl, 1 mM EDTA, 0.2% NP-40) resuspended in 2x SDS sample buffer, and denatured for 5 min at 95^°^C. Where indicated, total RNA prepared from WT yeast was added to the first wash step, and samples were rotated at 4^°^C for 10 min. Samples were analyzed by Western blot.

### Metabolite extraction and quantification

Metabolites were extracted as previously described^37^. LC-MS/MS mass spectrometric analyses were performed on a Sciex QTRAP 6500+ mass spectrometer equipped with an electrospray ion (ESI) source using scheduled MRM in the positive and negative mode separately. The ESI source was used in both positive and negative ion modes, configured as follows: Ion Source Gas 1 (Gas 1), 40psi; Ion Source Gas 2 (Gas 2), 45 psi; curtain gas (CUR), 45 psi; source temperature, 550°C; and ion spray voltage (IS), +5500 V(+) and −4500 V (−). The mass spectrometer was coupled to a Shimadzu HPLC (Nexera X2 LC-30AD). The system is controlled by Analyst 1.7.2 software.

Chromatography was performed under reversed-phase condition using an ACE 3 C18-PFP 150 x 4.6 mm HPLC column (Mac-Mod, USA). The column temperature, sample injection volume, the flow rate was set to 30°C, 5 μL, and 0.5 mL/min respectively. The HPLC conditions were as follows: Solvent A: Water with 0.1% Formic Acid (v/v), LC/MS grade and Solvent B: Acetonitrile with 0.1% Formic Acid (v/v), LC/MS grade. Gradient condition was 0-2 min: 5% B, 5-16 min: 90% B, 17 min 5% B, 30 min: 5% B. Total run time: 30 mins.

Negatively charged nucleotide metabolites were detected using TBA in negative mode as described previously^14^. Quantification was carried out using MultiQuant software by calculating peak area. Data was normalized to total ion count. Metabolite data are available in Table S1.

### RNA sequencing and analysis

RNA-seq was performed by Novogene. Library preparation was strand-specific and used rRNA removal. Strand-specific raw fastq reads were first evaluated using FastQC (https://www.bioinformatics.babraham.ac.uk/projects/fastqc/) and trimmed using TrimGalore (https://www.bioinformatics.babraham.ac.uk/projects/trim_galore/) with minimum Phred score of 35. To map paired-end reads we used STAR^38^ (v2.7.3a) with preset parameter value ranges suited for optimal alignment of reads to the yeast reference genome (S288C). Before mapping we first generated genome indices using both reference genome and reference annotations (R64-1-1) to get a more accurate alignment of the reads. Using Sambamba^39^ (v0.6.6) duplicates were marked, and flag statistics obtained with flagstat option. Uniquely mapped reads were quantified using featureCounts^40^ from Subread package (v1.6.3) per gene ID. To remove marked duplicates, “-- ignoreDup” was used and set “-s” to 1 for strand-specific read counting. In differential gene expression analysis fold changes and adjusted p-values were calculated from raw counts using the R package DESeq2^41^ (v3.16). Using obtained log fold changes, we defined up-regulated and down-regulated genes (with cutoff values of absolute fold change greater than or less than 1.5 and p_adj_ less than 0.05).For each gene set we then performed KEGG enrichment analysis using clusterProfiler package^42^. Dubious ORFs listed in the Saccharomyces Genome Database (SGD, https://www.yeastgenome.org/) were removed from entire datasets. All statistical analyses were performed in RStudio (v2022.07.2+576) and GraphPad Prism (v9.4.1). Raw sequencing data have been deposited at Gene Expression Omnibus with accession number: GSE #226779.

## Acknowledgements

We thank Dr. Hamid Baniasadi for assistance with the AB SCIEX QTRAP 6500^+^. We thank members of the Tu lab for helpful discussions. Some figures created using BioRender. This work was supported by grants from NIH R35GM136370, Welch Foundation I-1797, and investigator support from HHMI.

## Author Contributions

Conceptualization, M.M.M. and B.P.T.; Methodology, M.M.M. and B.M.S.; Investigation, M.M.M. and B.M.S.; Data analysis: M.M.M. and S.J.; Writing - Original Draft, M.M.M. and B.P.T.; Writing -Review & Editing, B.P.T.; Funding Acquisition, B.P.T.; Supervision, B.P.T.

## Ethics Declarations

The authors declare no competing interests.

## Data availability

All data generated or analyzed during this study are available in the main text or the Supplementary Information. Metabolite data are available in Supplementary Table S1. RNA-seq data are available in Supplementary Table S2. Sequencing data have been deposited to GEO with accession number GSE #226779.

---

**Fig. S1.**
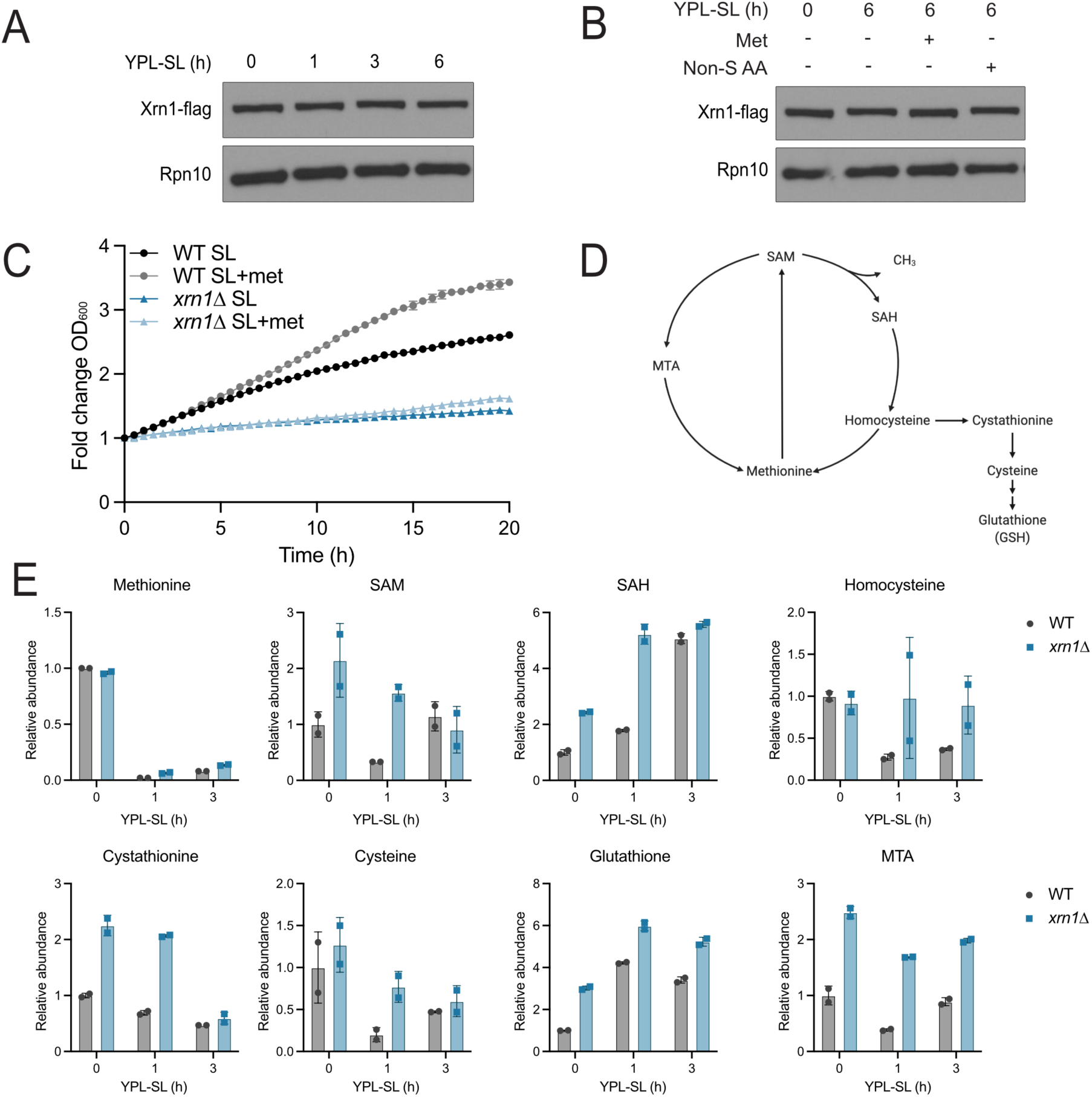
Loss of Xrn1 causes elevated levels of SAM and other sulfur-containing metabolites. **(A)**, **(B)** Xrn1 protein abundance is not altered under different metabolic conditions. Anti-flag Western blot assessing protein amounts of Xrn1 under the indicated conditions. **(C)** Methionine restores growth of WT but not *xrn1Δ* cells. Growth curve measuring OD_600_ of WT or *xrn1Δ* cells in the indicated media. OD_600_ was measured every 30 min. Fold change is plotted. The data are represented as mean ± SD (n=3). **(D)** Schematic of sulfur-containing metabolites in yeast produced from methionine and transsulfuration. **(E)** Many sulfur-containing metabolites are elevated in cells lacking Xrn1. WT and *xrn1Δ* cells were grown in the indicated conditions. Metabolite samples were collected and analyzed by LC-MS/MS. Data are represented as mean ± SD (n=2). The metabolite data are also presented in Table S1.

**Fig. S2.**
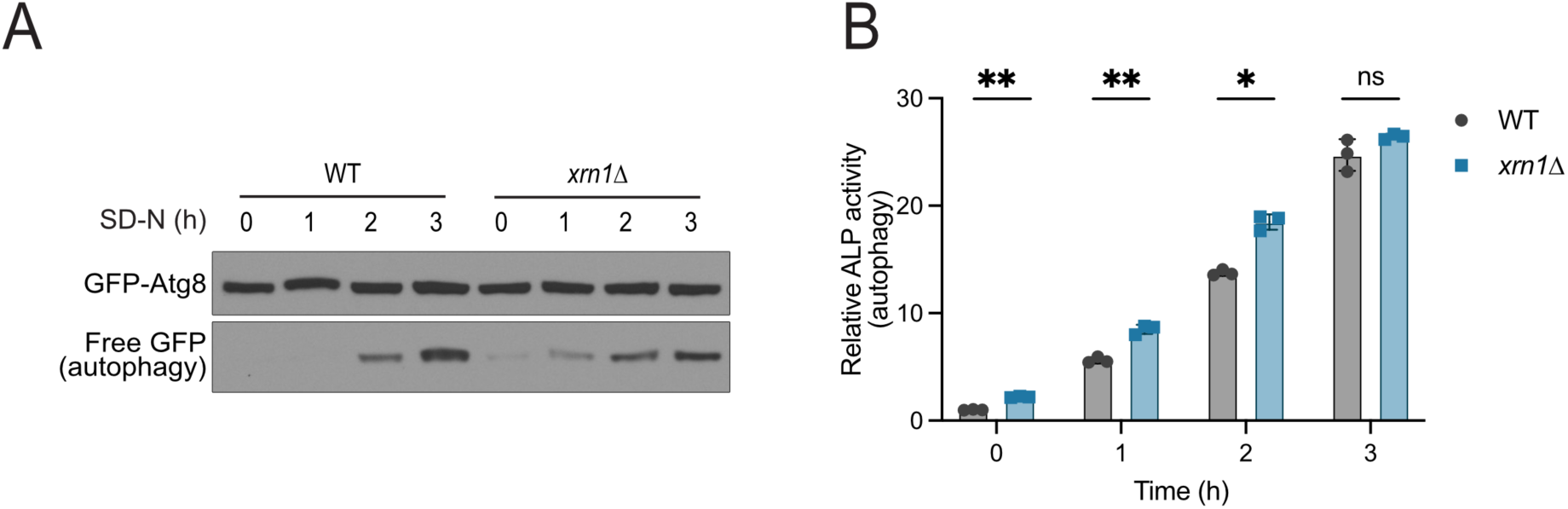
Xrn1 is a negative regulator of nitrogen starvation-induced autophagy. **(A)** Autophagy under nitrogen starvation conditions is induced more rapidly in *xrn1Δ* cells. WT and *xrn1Δ* cells harboring a centromeric plasmid expressing GFP-Atg8 were grown to mid-log phase in YPD then starved for nitrogen (SD-N) for the indicated times. Free GFP, indicative of autophagy induction, was detected by Western blot. **(B)** Autophagy under nitrogen starvation conditions in *xrn1Δ* cells as monitored by ALP assay. Cells were grown to mid-log phase in YPD then starved of nitrogen (SD-N) for the indicated times. ALP activity was measured and normalized to the WT cells in rich media. Mean±SD, n=3, statistical analysis performed using student’s t-test.

**Fig. S3.**
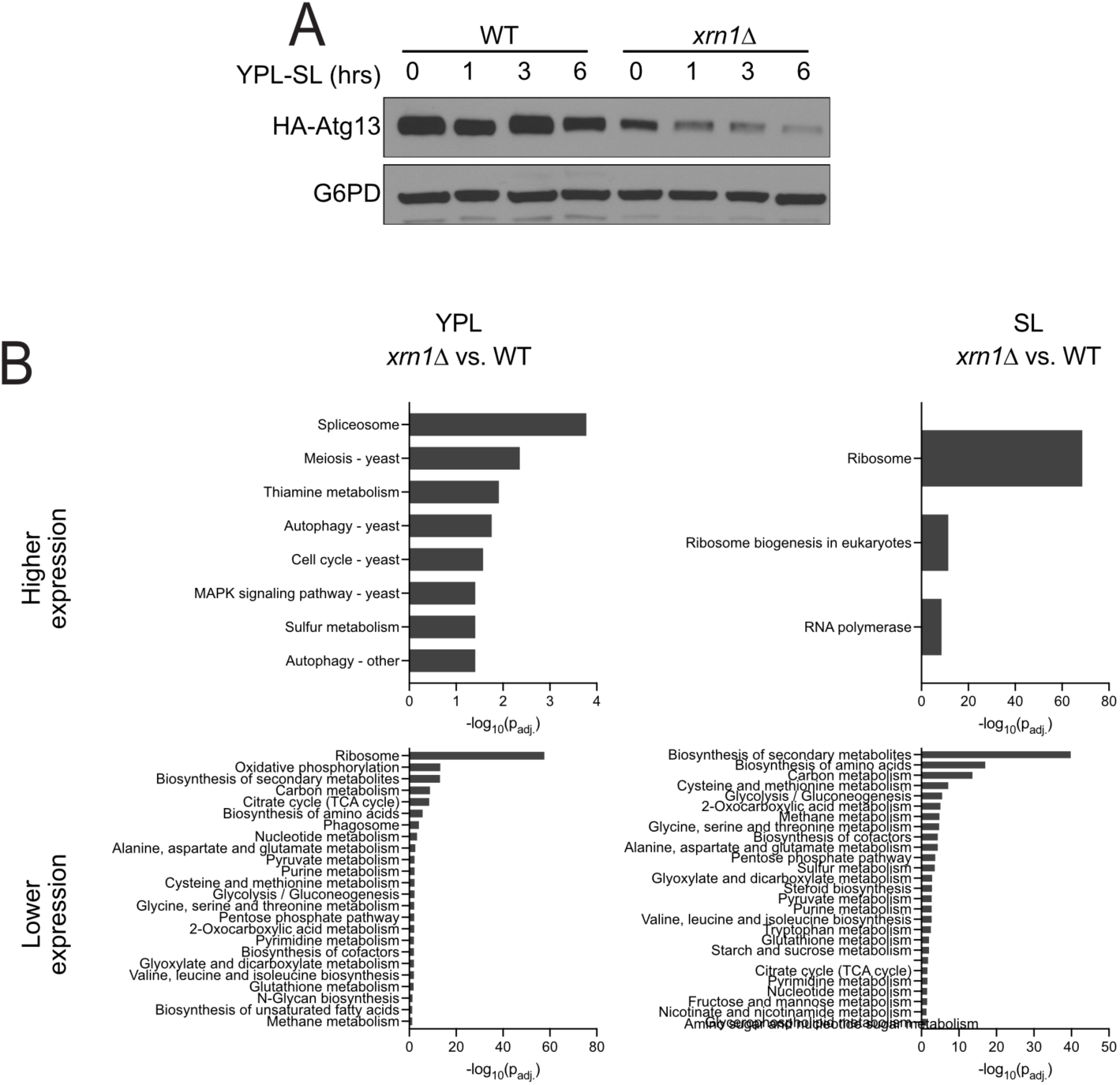
Amounts of select autophagy mRNAs and proteins in cells lacking Xrn1. **(A)** Key autophagy protein Atg13 exhibits reduced abundance in cells lacking Xrn1. HA-tagged Atg13 protein abundance was assayed in WT and *xrn1Δ* cells following methionine deprivation for the indicated times by Western blotting. **(B)** Gene ontology enrichment from RNA-seq identifies gene groups that are altered in *xrn1Δ* cells.

**Fig. S4.**
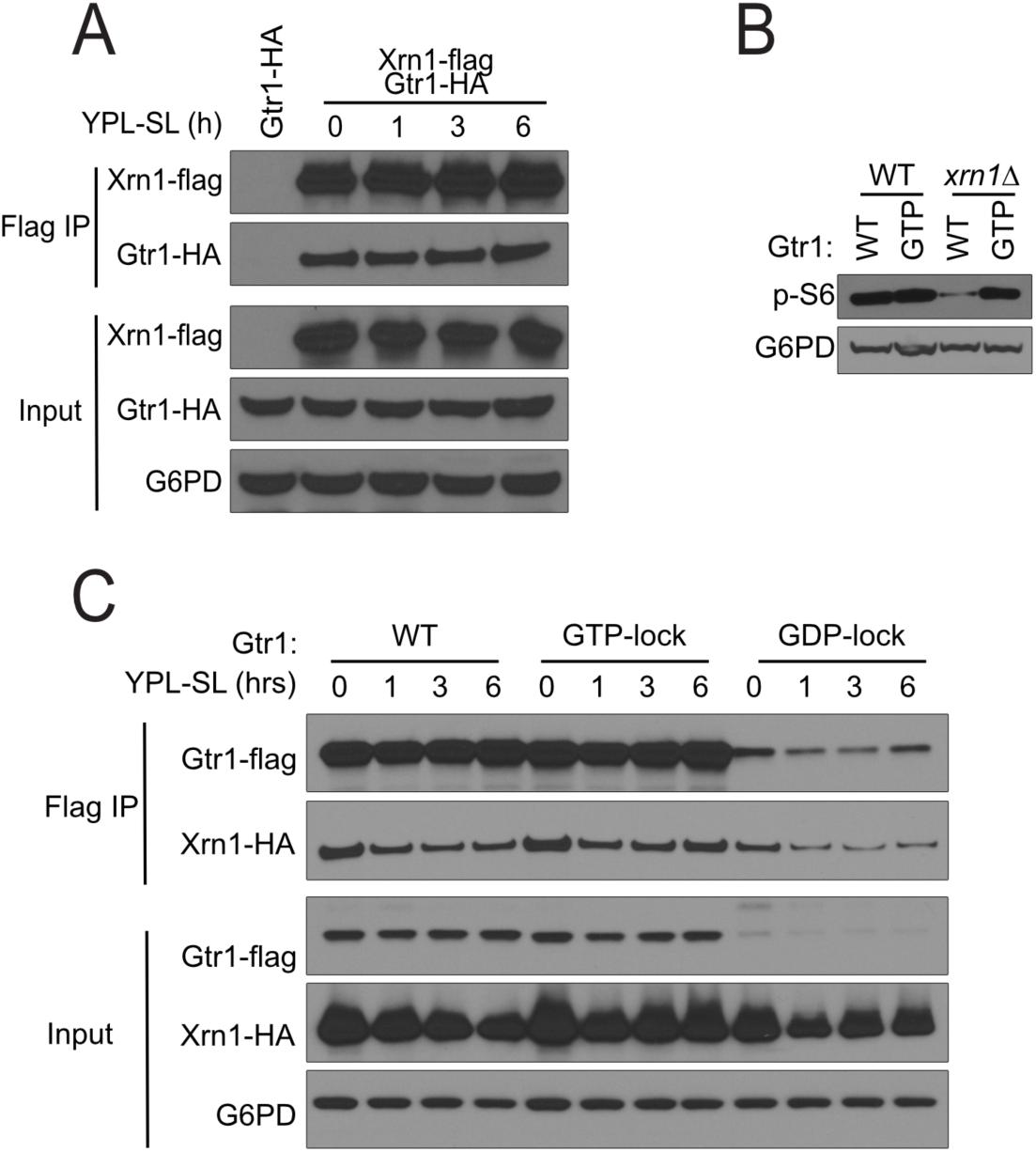
Xrn1 interacts with Rag GTPase Gtr1. **(A)** Xrn1 and Gtr1 interact independent of methionine availability. Cells expressing flag-tagged Xrn1 and HA-tagged Gtr1 were grown in YPL and switched to SL for the indicated times. Interaction between these proteins was assessed by co-IP followed by Western blot. **(B)** GTP-locked mutation of Gtr1 restores TORC1 activity in cells lacking *xrn1Δ*, assayed by Western blot for phosphorylated S6 ribosomal protein. **(C)** Xrn1 preferentially interacts with the GDP-locked form of Gtr1 by co-IP. Cells expressing flag-tagged Gtr1 constructs and HA-tagged Xrn1 were grown in the indicated conditions. Interaction between these proteins was assessed by co-IP followed by Western blot. Note the GDP-locked (S20L) mutation destabilizes the Gtr1 protein compared to WT or GTP-locked (Q65L) mutation.

**Fig. S5.**
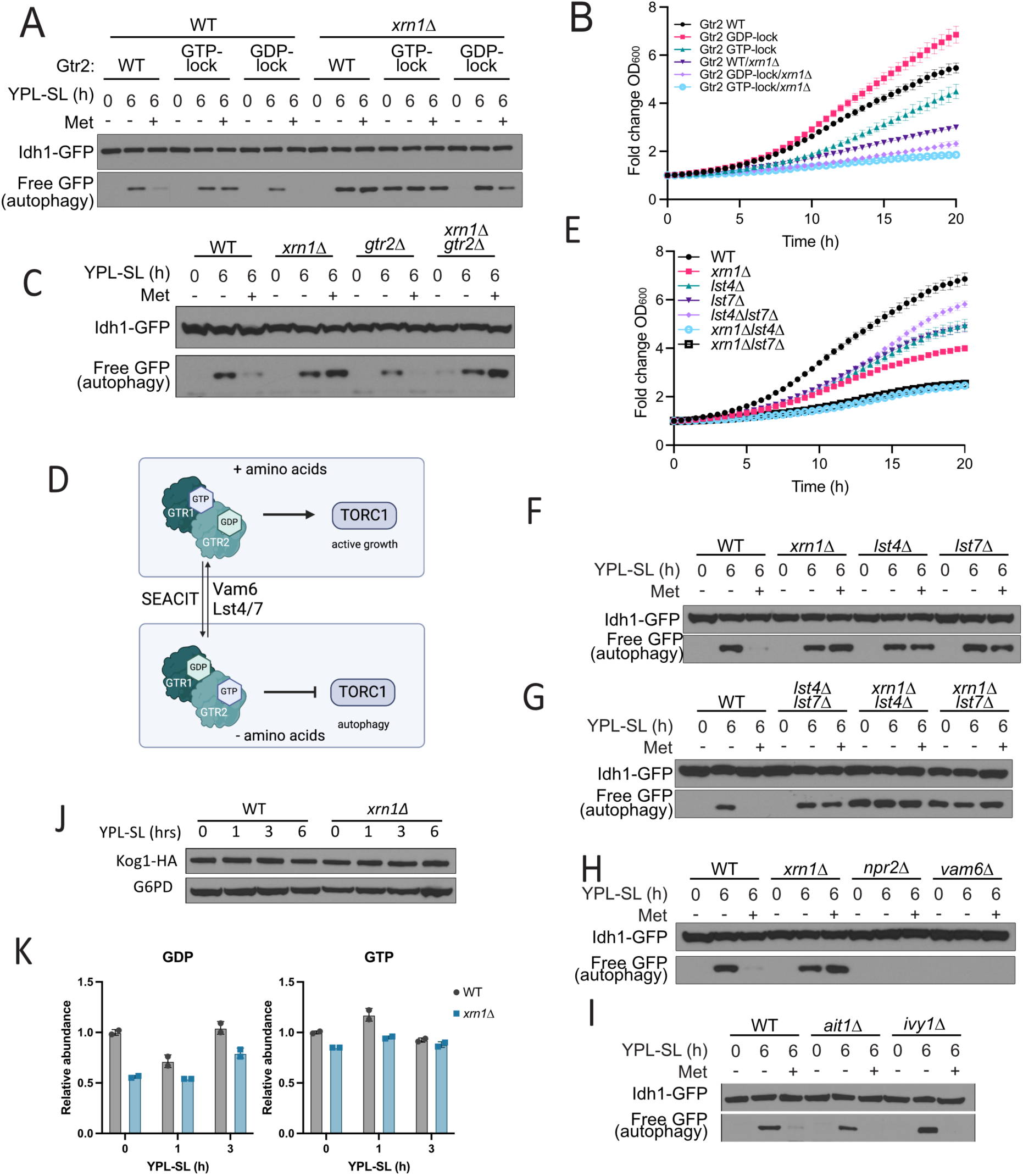
Xrn1 does not act through known Gtr1/Gtr2 regulatory proteins. **A)** GTP-and GDP-locked mutations in Gtr2 synergize with loss of Xrn1 in regulating autophagy following methionine deprivation. The indicated strains expressing point mutations of Gtr2 to lock it in its GTP-binding (Q66L) or GDP-binding (S23L) state were grown in YPL and then switched to SL for 6 h, in the absence or presence of 1 mM methionine. Autophagy was measured by the GFP cleavage assay. **(B)** GTP-and GDP-locked mutations in Gtr2 exacerbate the growth defect of *xrn1Δ* cells. The indicated strains were grown in YPL and OD_600_ was measured every 30 min. Fold change in OD_600_ is plotted. **(C)** Cells lacking both Xrn1 and Gtr2 exhibit enhanced autophagy. The indicated strains were assayed for autophagy as in (A). **(D)** Schematic depicting how the nucleotide binding states of Gtr1 and Gtr2 are controlled by GAPs and GEFs. **(E)** Growth curves of cells lacking Xrn1 and either Lst4 or Lst7 reveals a synthetic growth defect. The indicated strains were grown in YPL and OD_600_ was measured every 30 min. Fold change in OD_600_ is plotted. **(F)** Loss of the Gtr2 GAPs Lst4 and Lst7 phenocopy the methionine-insensitive autophagy phenotype of *xrn1Δ* mutants. Autophagy was assayed as in (A). **(G)** Cells lacking Xrn1 in combination with Lst4 or Lst7 exhibit significantly enhanced autophagy. Autophagy was assayed as described as in (A). **(H)** Loss of Vam6 phenocopies *npr2Δ* cells in regulation of autophagy. Vam6 is annotated as a GEF for Gtr1. The indicated strains were assayed for autophagy as in (A). **(I)** Loss of Ait1 and Ivy1 do not play a role in regulation of autophagy in response to methionine deprivation. The indicated strains were assayed for autophagy as in (A). **(J)** Abundance of Kog1 protein is not altered in *xrn1Δ* compared to WT. Either WT or *xrn1Δ* with HA-tagged Kog1 were grown in the indicated conditions, and Kog1 protein abundance was assayed by Western blotting. **(K)** Targeted metabolomics reveals slightly reduced levels of both GDP and GTP in cells lacking Xrn1. WT and *xrn1Δ* cells were grown in the indicated conditions. Metabolites were extracted at the indicated times and analyzed by targeted LC-MS/MS.

**Table S1:**
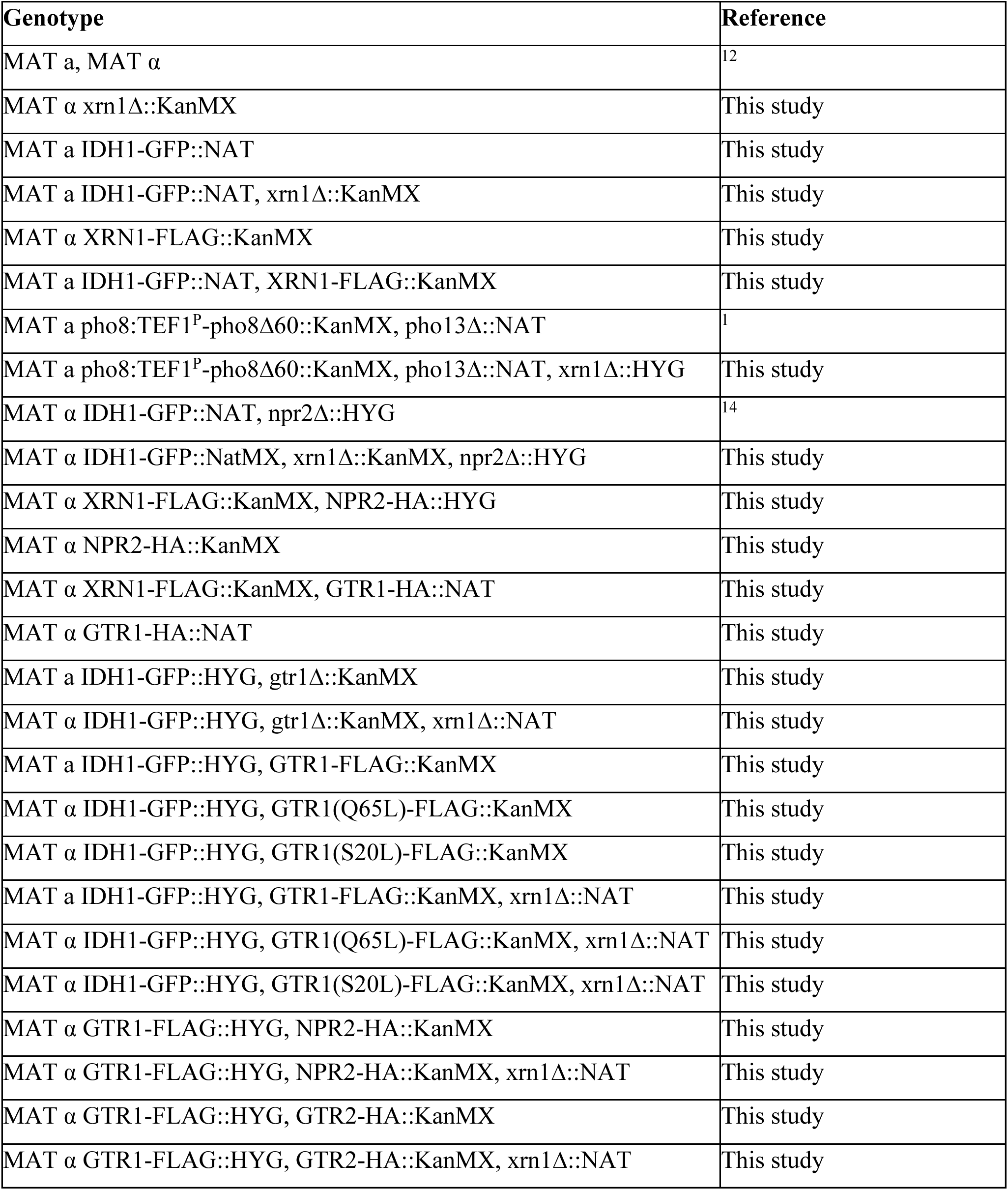
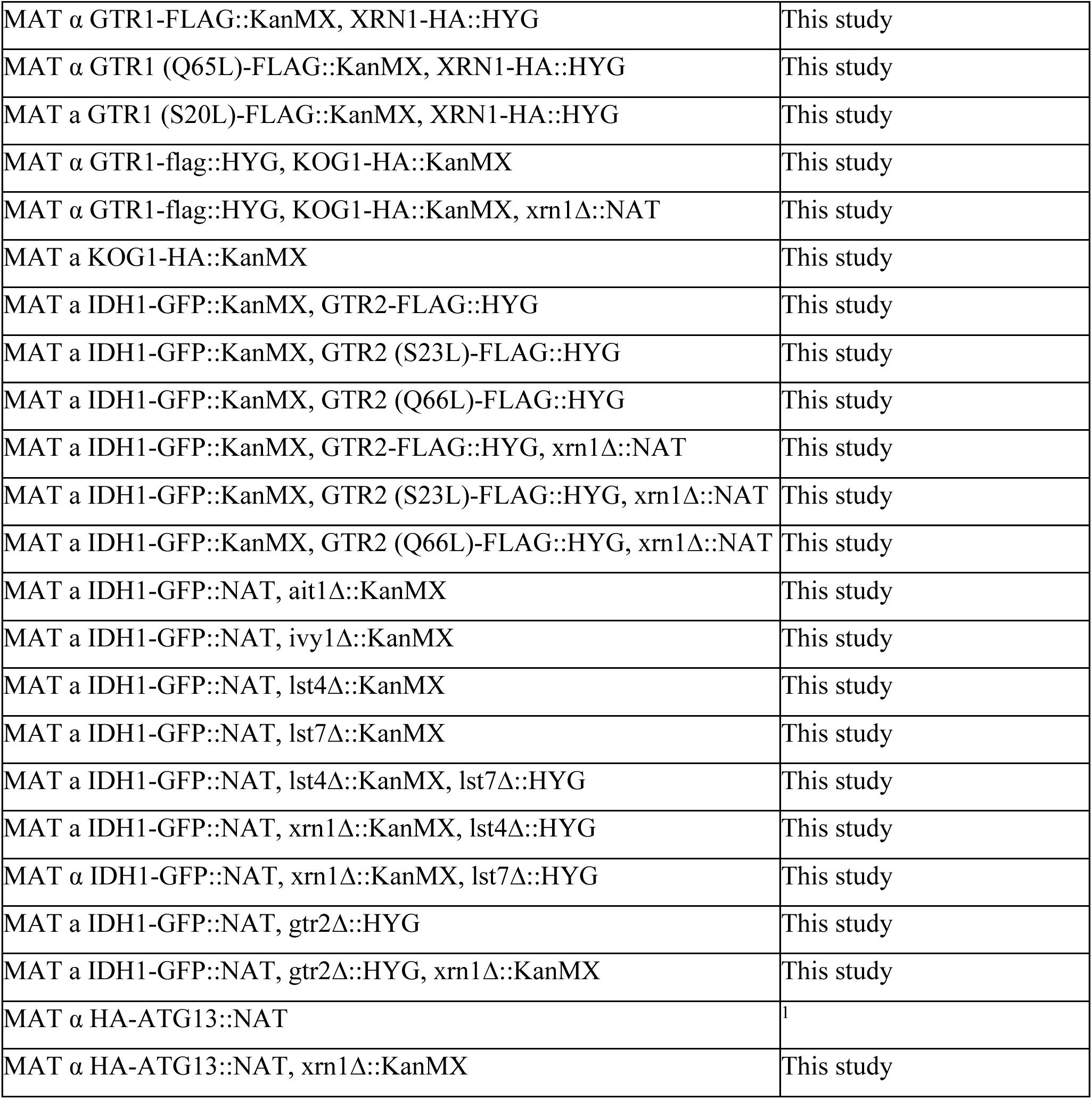
Yeast strains used in this study. All strains are in the prototrophic CEN.PK background.

